# Single-cell transcriptional profiling in Arabidopsis root exposed to B toxicity at seedling stages

**DOI:** 10.1101/2023.03.09.531923

**Authors:** Hikmet Yılmaz, Ceyhun Kayıhan, Halis Batuhan Ünal, Oğuzhan Yaprak, Emre Aksoy

## Abstract

Cell-specific transcriptional responses to environmental stimuli are yet to be fully characterized in plants. In this study, we apply single-cell RNA sequencing to *Arabidopsis thaliana* roots exposed to boron (B) toxicity to characterize the transcription map at cellular resolution and thus, to understand how B toxicity can alter gene expression and development at single cell resolution. Single-cell transcriptomes from protoplasts of more than 2750 *Arabidopsis thaliana* root cells were obtained. Plotting the single-cell transcriptomes via t-SNE projections yielded six major cell clusters including quiescent cells (QC), endodermis, cortex, columella, trichoblast (root-hair), and root cap. The maximum number of most significantly upregulated genes were determined in columella under 1 mM B and in endodermis under 2 mM B condition. Additionally, the maximum number of most significantly upregulated genes under 3 mM B and 5 mM B conditions was determined in the root cap, implying a critical role against severe B toxicity conditions. We also showed that these upregulated genes are highly correlated with “glutathione metabolism” in columella and “carbon metabolism” in root cap. Taken together, for the first time in the literature, our study provides a gene expression map at single-cell resolution and describes the extent of heterogeneity at the molecular level among populations of different cell types in Arabidopsis root under B toxicity conditions.

## Introduction

B is an essential micronutrient for plant development and growth. Because of the poor drainage in arid and semiarid countries such as Morocco, Syria, Egypt, Iraq, Italy, and Turkey, B can easily accumulate in the soil (Nable et al., 1997; Pennisi et al., 2006). When its concentration is slightly higher than necessary for growth in plants, it can become toxic and severely affect plant growth and cause yield losses in crop production (Mengel & Kirkby, 2012) because B accumulation at a toxic level in the soil leads to impairment of growth and plant metabolism, causing chlorosis and necrosis in leaf tissues (Reid et al., 2004; Landi et al., 2012; Camacho-Cristóbal et al., 2018). B can bind to the cis-hydroxyl groups of biologically important molecules including cis-hydroxyl groups such as ribose, sorbitol, and apiose and thereby it can form strong complexes with them. For this reason, toxic amount of B cause metabolic damage to plant cells by binding to ribose-containing biomolecules such as ATP, nicotinamide adenine dinucleotide and nicotinamide adenine dinucleotide phosphate and by binding to ribose in RNA, it may cause disruption of cell division and disruption of cell growth (Loomis & Durst, 1992; Reid, et al., 2004). Thus, B toxicity may cause severe physiological disorders including photosynthesis inhibition (Lovatt & Bates, 1984), increase in membrane leakage and lipid peroxidation (Karabal, et al., 2003) and change in antioxidant enzyme activity (Karabal, et al., 2003) due to overaccumulation of reactive oxygen species (ROS) caused by oxidative stress.

B-tolerant cultivars are characterized by decreased B concentration in their leaf tissues under toxic B conditions (Nable et al. 1990). However, it has been suggested that the high B accumulation in shoot tissue does not always correlate with the severity of B toxicity symptoms (Nable et al., 1997 and Sakamoto et al., 2011). Reduced uptake of B by roots in some genotypes has an important role in improved B tolerance (Torun et al. 2006). Likewise, the physiological basis for reduced B accumulation in tolerant cultivars was shown to be related to the active efflux of B from root cells (Hayes and Reid 2004). Besides root uptake, root-to-shoot transport, shoot accumulation of B, and internal mechanisms at cellular level are involved in detoxification of B, and thereby, B tolerance in some genotypes (Torun et al. 2006). However, it is clear that much is unknown about B regulation mechanisms. Therefore, identifying genes that contribute to B-stress response and characterization of these genes are critical to improve tolerance of plants to B toxicity. Although limited number of studies focusing on toxic B responsive mechanisms at molecular level have been conducted in plants, significant contributions have been made in improving our understanding of the effects of B stresses and the tolerance mechanism (Feng, et al., 202; Kayıhan C., 2021; Chen, et al., 2022; Pandey, et al., 2022).

Also, differentially regulated genes in response to B toxicity have been identified and various pathways have been found in various plants by using bulk methods such as microarray and RNA-sequencing (Öz, et al., 2009; Aquea, et al., 2012; Ozhuner, et al., 2013; Huang, et al., 2016; Kayıhan, et al., 2017; deAbreu-Neto, et al., 2017; Kayıhan, et al., 2019; Huang, et al., 2019; Brdar-Jokanović, 2020). Accordingly, it has been mainly found that WRKY, MYB, and NAC transcription factors have been proposed to be involved in regulation of B homeostasis in plants. However, these bulk methods can yield qualitatively inaccurate results because different plant cell populations are averaged (Trapnell, et al., 2014).

A recently developed single-cell RNA sequencing (scRNA-seq) profiles the transcriptome of hundreds to thousands of individual cells, and thus enables us to understand the cell as a functional unit and to gain insight into tissue cellular heterogeneity (Macosko, et al., 2015; Ziegenhain, et al., 2017). It allows us to identify subpopulations of a known cell type using differences in gene expression patterns within the cell population. It may also identify previously unknown cell types (Artegiani, et al., 2017; Villani, et al., 2017; Glass, et al., 2017). In addition, scRNA-seq gives “ trajectories ” of the differentiation process in cell populations with differentiation at different time points and each differentiation status of cells from these trajectories (Treutlein, et al., 2014; Trapnell, et al., 2014; Artegiani, et al., 2017; Qiu, et al., 2017). Although scRNA-seq is not widely applied in plants compared to animals, several groups have recently applied the efficient scRNA-seq to plants as Arabidopsis and rice. Especially, Arabidopsis root provides a useful organ for the application of scRNA-seq due to having small number of cells and cell types and availability of isolating individual cells via protoplasting and, until now, many cell-type marker genes have been identified in these plants (Bruex, et al., 2012; Birnbaum, et al., 2015; Efroni, et al., 2015; Li, et al., 2016; Ryu, et al., 2019; Wendrich, et al., 2020; Wang, et al., 2021; Apelt, et al., 2022).

In this study, transcriptomic profiles in individual protoplasts from Arabidopsis roots exposed to B toxicity at seedling stages were determined by using a commercially available droplet-based platform (10X Genomics Chromium; Zheng, et al., 2017) to analyze a gene expression map of Arabidopsis root in response to B toxicity at single-cell resolution. Accordingly, cell clusters related to quiescent center (QC), endodermis, root cap, columella, cortex, and trichoblast have been identified in this single-cell transcriptome. Trichoblast was not identified in Arabidopsis root under B toxicity conditions. Also, same genes related to B tolerance mechanisms previously presented in the literature have been detected, many new genes have been identified at specific cell types. Finally, specific genes at these cell clusters might be used as markers, and this can affect transgenic and breeding strategies for B tolerance mechanism in plants.

## Results

### scRNA-seq and identification of cell type clusters

Isolated protoplasts from Arabidopsis roots exposed to B toxicity at seedling stages were used by using the 10X Genomics Chromium platform (Zheng, et al., 2017) to generate a high-resolution scRNA-seq. In summary, a total population of 2755 cells were recovered across two replicates. Approximately 304,322 reads were obtained per cell, which generated a median of 1,957 unique molecular identifiers per cell, more than 15000 total genes detected in the population, and more than 93% valid barcode total per each replicate (Supplemental Dataset S1). Plotting the single-cell transcriptomes via Louvain clustering and t-SNE projections using Loupe Browser (v.6.2.0). yielded six clusters of cell transcriptomes. We then performed the tissue/cell type assignment to clusters in our single-cell population by using fourteen root tissue/cell types specific marker genes (Supplemental Dataset S2-3). They led to the assignment of specific cell clusters to quiescent cells (QC), endodermis, cortex, columella, trichoblast (root-hair) and root cap under control and endodermis, QC, root cap and columella B toxicity conditions (Figure 1). Cortex and trichoblast cell clusters were not detected in Arabidopsis root exposed to all B toxicity treatments.

**Figure 1.**
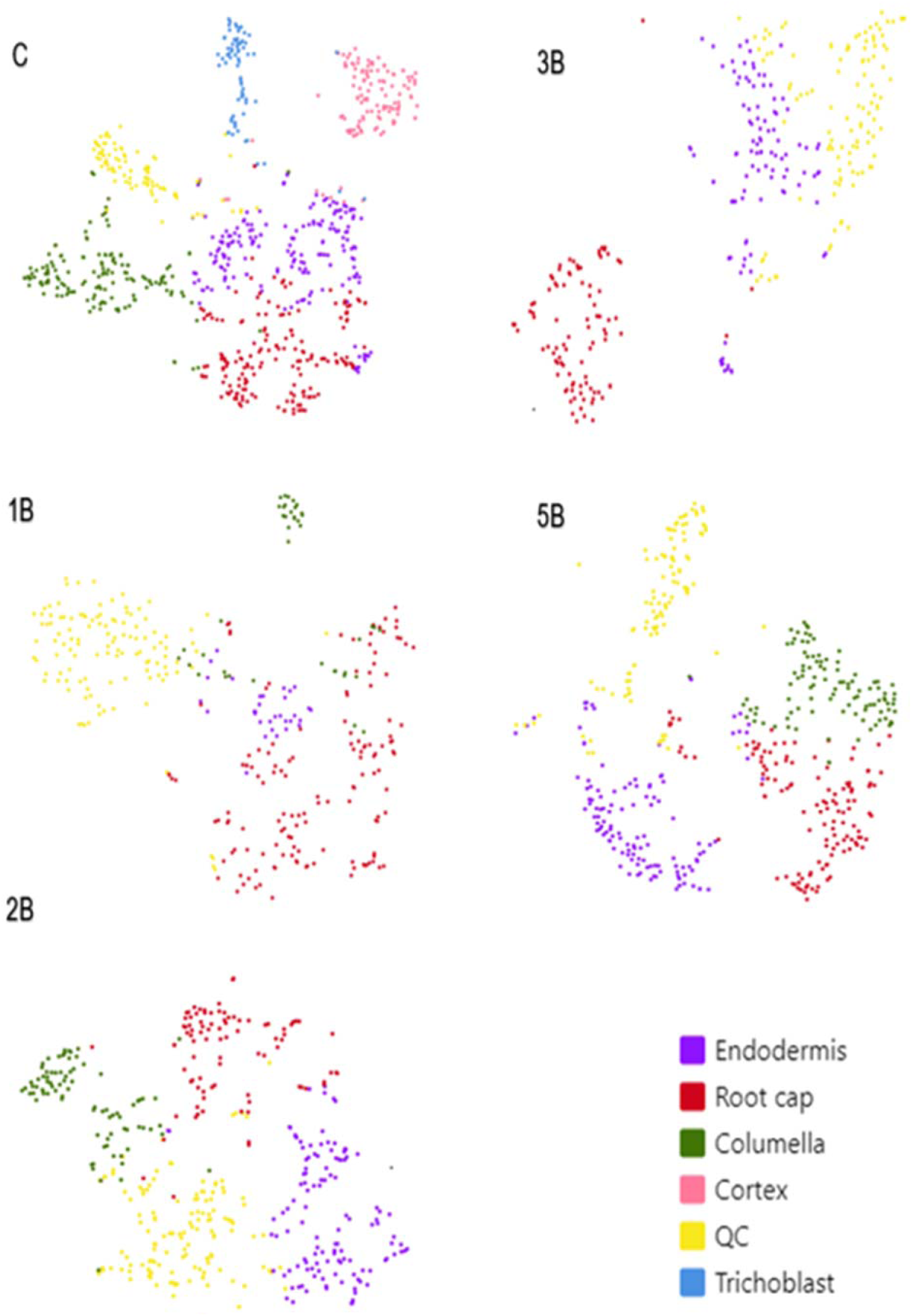
t-SNE visualization to identify putative 6 cell clusters from Arabidopsis roots. Each dot denotes a single cell. Colors denote corresponding cell clusters. (QC: Quiescent center). C: Control, 1B: 1 mM boric acid treatment, 2B: 2 mM boric acid treatment, 3B: 3 mM boric acid treatment and 5B: 5 mM boric acid treatment.

### Differentially expressed Genes (DEGs) of single-cell transcriptome of Arabidopsis roots exposed to B toxicity

We used Loupe software (v.6.2.0) to identify changes in gene expression profiles among all clusters (Supplemental Dataset S4) and for each cluster individually (Supplemental Dataset S5-9) in Arabidopsis roots under B toxicity. The number of overlaps DEGs between all group were shown in Figure 2, A-B, and the number of overlaps DEGs in the clusters for each group were shown in Figure 2, C-L, and the number of overlaps DEGs between the groups for each cluster were shown in Figure 3 by using upset plot (http://www.bioinformatics.com.cn/en). Additionally, the overlap DEGs between the clusters for each group and the overlap DEGs between the groups for each cluster were shown in Supplemental Dataset S10-11 by using Venn diagram (https://bioinformatics.psb.ugent.be/webtools/Venn/). Accordingly, we found that 71, 34, 165, 32 and 147 genes were specifically upregulated in C, 1B, 2B, 3B and 5B, respectively (Figure 2A; Supplemental Dataset S10). 51 and 11 genes were commonly upregulated between 2B and 3B, and between 1B and 5B, respectively and 16 genes were commonly upregulated in 1B, 2B and 3B (Figure 2A; Supplemental Dataset S10). 236, 16, 93, 79 and 35 genes were specifically downregulated in C, 1B, 2B, 3B and 5B, respectively (Figure 2B; Supplemental Dataset S10). 29 genes were commonly downregulated between 1B and 5B (Figure 2B; Supplemental Dataset S10). 7 genes were commonly downregulated in C, 1B, 2B, 3B and 5B, and 7 genes were commonly downregulated in 1B, 2B, 3B and 5B (Figure 2B; Supplemental Dataset S10).

**Figure 2.**
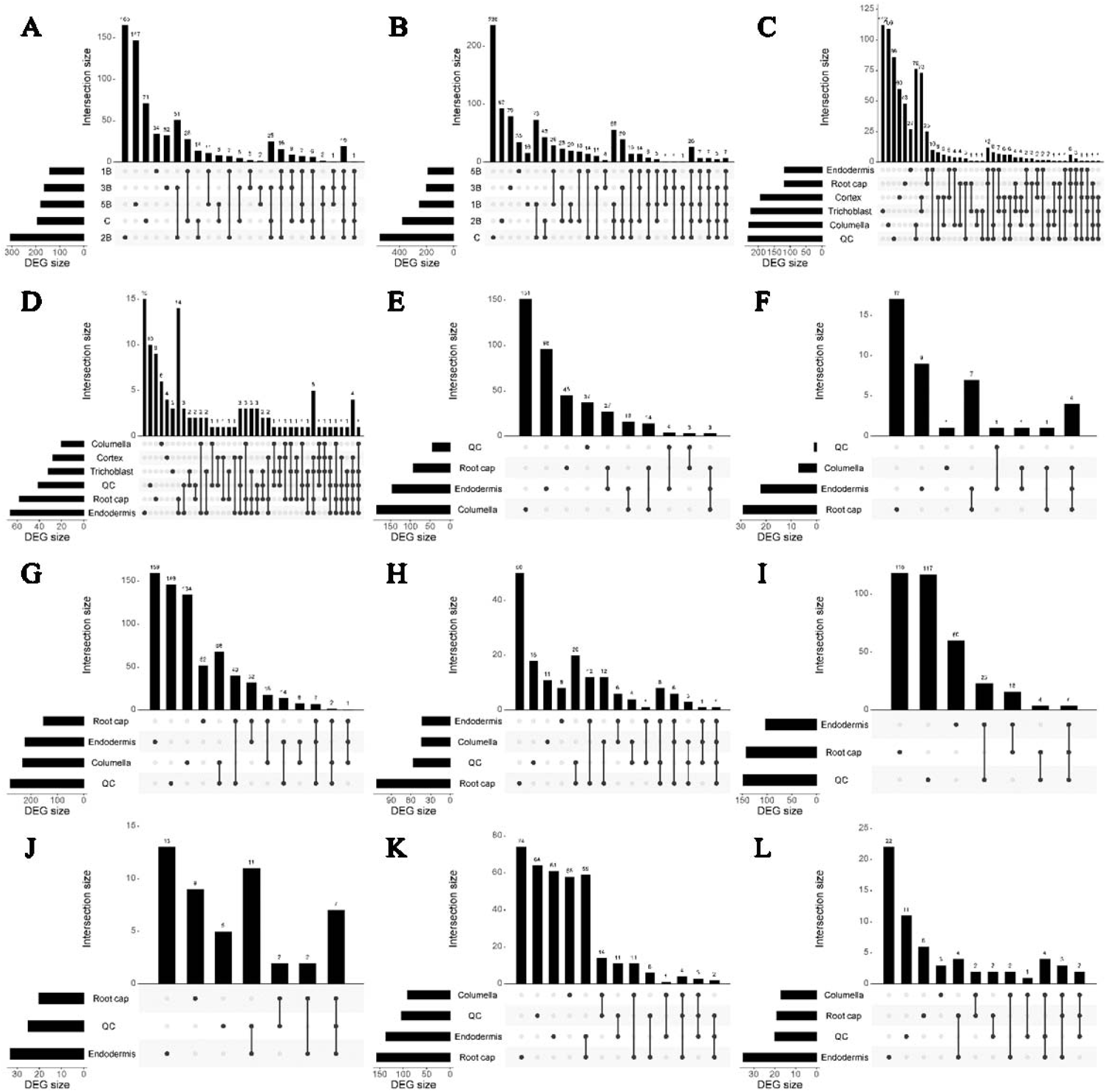
Upset plots to summarize the common and DEGs under control (C) and B toxicity treatments in root tissues of Arabidopsis thaliana. 1B: 1 mM boric acid treatment, 2B: 2 mM boric acid treatment, 3B: 3 mM boric acid treatment and 5B: 5 mM boric acid treatment, (A): the intersection of upregulated genes between C and B toxicity treatment groups, (B): the intersection of downregulated genes between C and B toxicity treatment groups, (C): the intersection of upregulated genes between clusters under C condition, (D): the intersection of downregulated genes between clusters under C condition (E): the intersection of upregulated genes between clusters under 1B condition, (F): the intersection of downregulated genes between clusters under 1B condition, (G): the intersection of upregulated genes between clusters under 2B condition, (H): the intersection of downregulated genes between clusters under 2B condition, (I): the intersection of upregulated genes between clusters under 3B condition, (J): the intersection of downregulated genes between clusters under 3B condition, (K): the intersection of upregulated genes between clusters under 5B condition and (L): the intersection of downregulated genes between clusters under 5B condition. In each panel, the bottom left horizontal bar graph labeled DEG size shows the total number of DEGs per post-treatment time point. The circles in each panel’s matrix represent what would be the different Venn diagram sections (unique and overlapping DEGs.

**Figure 3.**
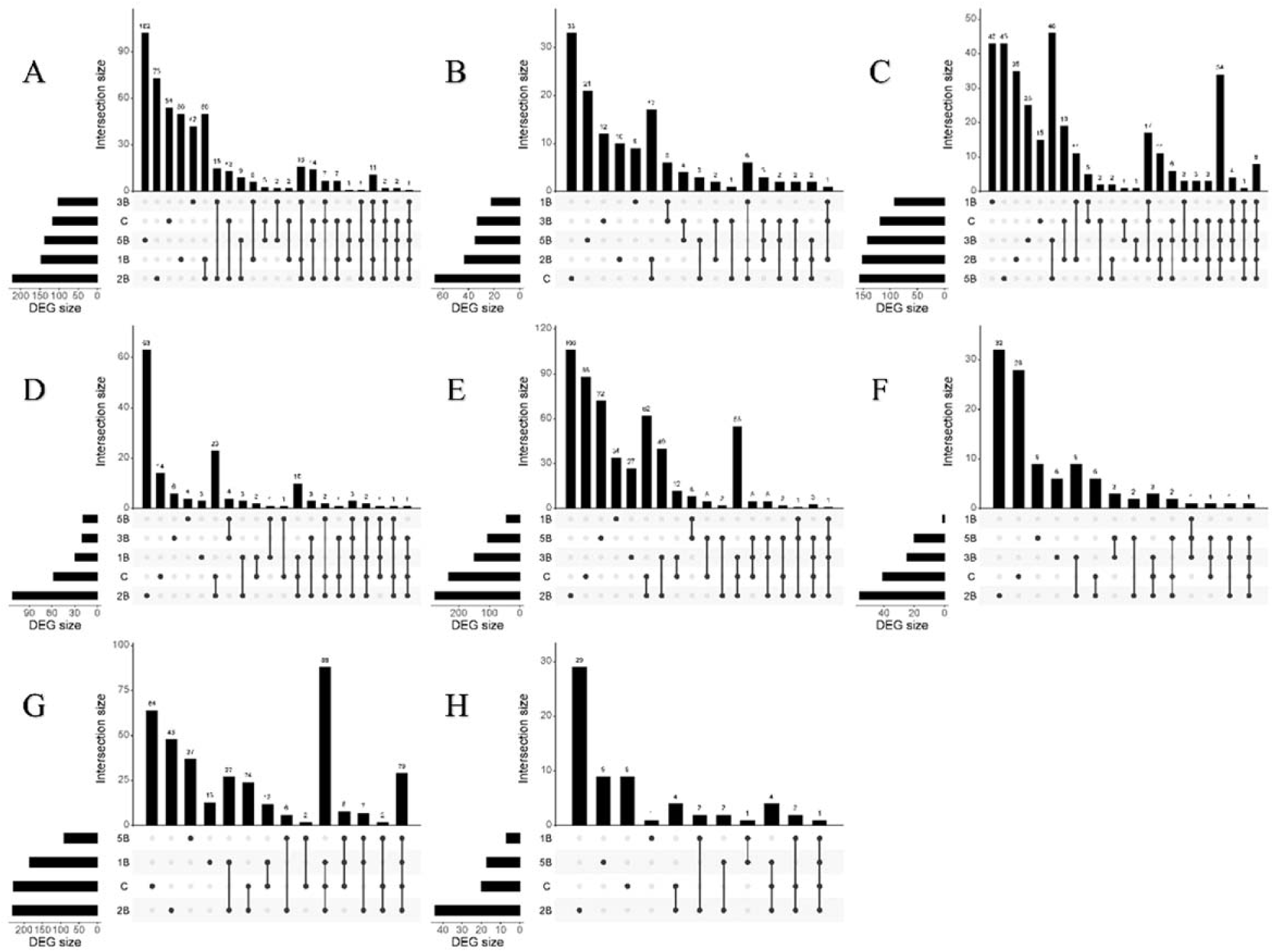
Upset plots to summarize overlap between clusters for up- or down-regulated. A, C, E, G) the intersection up-regulated genes in Endodermis, Root cap, QC and Columella respectively, and B, D, F, H) the intersection down-regulated genes in Endodermis, Root cap, QC and Columella, respectively. In each panel, the bottom left horizontal bar graph labeled DEG size shows the total number of DEGs per post-treatment time point. The circles in each panel’s matrix represent what would be the different Venn diagram sections (unique and overlapping DEGs.

We also determined the overlapping DEGs between clusters for each group (Figure 2, C-L; Supplemental Dataset S10). Accordingly, we found that 112, 109, 86, 60, 48 and 27 genes were specifically upregulated under C in trichoblast, columella, QC, cortex, root cap and endodermis, respectively (Figure 2C; Supplemental Dataset S10). Moreover, under this condition, 76, 73, 25 and 10 genes were commonly upregulated between columella and QC, and between cortex and trichoblast, and between endodermis and root cap, and between endodermis and QC, respectively, and also 12 genes were commonly upregulated in endodermis, root cap and QC and 8 genes were commonly upregulated in endodermis, columella and QC, and 7 genes were commonly upregulated in endodermis, cortex and trichoblast (Figure 2C; Supplemental Dataset S10). 15, 10, 9, 6, 4 and 3 genes were specifically downregulated under C in endodermis, QC, root cap, columella, cortex, and trichoblast, respectively (Figure 2D; Supplemental Dataset S10). Furthermore, 14 genes were commonly downregulated between endodermis and root cap and also, 5 genes were commonly downregulated in cortex, trichoblast, root cap and endodermis, and 4 genes were commonly downregulated in cortex, trichoblast, QC, root cap and endodermis (Figure 2D; Supplemental Dataset S10).

151, 96, 45 and 37 genes were specifically upregulated under 1B in columella, endodermis, root cap and QC, respectively (Figure 2E). Additionally, 27, 16 and 14 genes were commonly upregulated between root cap and endodermis, and between endodermis and columella, and between root cap and columella, respectively (Figure 2E; Supplemental Dataset S10). 17, 9 and 1 genes were specifically downregulated under 1B in root cap, endodermis and columella, respectively (Figure 2F; Supplemental Dataset S10). Moreover, 7 genes commonly downregulated between root cap and endodermis (Figure 2F; Supplemental Dataset S10). 159, 146, 134 and 52 genes were specifically upregulated under 2B in endodermis, QC, columella, and root cap, respectively (Figure 2G; Supplemental Dataset S10). Also, 68, 40, 32, 18 and 14 genes were commonly upregulated between columella and QC, and between root cap and QC, and between root cap and endodermis, and between root cap and columella, and between endodermis and QC, respectively. Furthermore, 7 genes commonly upregulated in root cap, endodermis and QC (Figure 2G; Supplemental Dataset S10).

50, 18, 11 and 8 genes were specifically downregulated under 2B in root cap, QC, columella and endodermis, respectively (Figure 2H; Supplemental Dataset S10). Additionally, 20, 12 and 12 genes were commonly downregulated between QC and root cap, and between endodermis and root cap, and between columellar and root cap, respectively. Also, 8 genes commonly downregulated in endodermis, QC and root cap, and 6 genes commonly downregulated in endodermis, columella and root cap (Figure 2H; Supplemental Dataset S10).

118, 117 and 60 genes were specifically upregulated under 3B in root cap, QC and endodermis, respectively (Figure 2I; Supplemental Dataset S10). Moreover, 23 and 16 genes were commonly upregulated between endodermis and QC, and between endodermis and root cap, respectively, and also 4 genes commonly upregulated in root cap, endodermis and QC (Figure 2I). 13 and 9 genes were specifically downregulated under 3B in endodermis and root cap, respectively (Figure 2J; Supplemental Dataset S10). Also, 11 genes were commonly downregulated between endodermis and QC, and 7 genes were commonly downregulated in endodermis, QC and root cap (Figure 2J; Supplemental Dataset S10).

74, 64, 61 and 58 genes were specifically upregulated under 5B in root cap, QC, endodermis and columella, respectively (Figure 2K; Supplemental Dataset S10). Additionally, 59, 14, 11 and 11 genes were commonly upregulated between endodermis and root cap, and between columella and QC, and between QC and endodermis, and between columella and root cap, respectively (Figure 2K; Supplemental Dataset S10). 22, 11, 6 and 3 genes were specifically downregulated under 5B in endodermis, QC, root cap and columella, respectively (Figure 2L; Supplemental Dataset S10). Moreover, 4 genes were commonly downregulated in columella, QC and endodermis (Figure 2L; Supplemental Dataset S10).

Furthermore, we determined the common and DEGs between C and B toxicity conditions for each cell cluster to find the high B responsive regulations of gene expression patterns of clusters in Arabidopsis root (Figure 3; Supplemental Dataset S11). Accordingly, 54, 50, 73, 42 and 102 genes were specifically upregulated in endodermis under C, 1B, 2B, 3B and 5B, respectively (Figure 3A; Supplemental Dataset S11). Moreover, in this cluster, 50, 15, 9 and 6 genes were commonly upregulated between 1B and 2B, and between 2B and 3B, and between 2B and 5B, and between 1B and 3B, respectively, and also 16 genes were commonly upregulated in 1B, 2B and 3B (Figure 3A; Supplemental Dataset S11). On the other hand, 33, 9, 10, 12 and 21 genes were specifically downregulated in endodermis under C, 1B, 2B, 3B and 5B, respectively (Figure 3B; Supplemental Dataset S11). Also, 6 and 4 genes were commonly downregulated between 1B and 3B, and between 3B and 5B, respectively (Figure 3B; Supplemental Dataset S11). 15, 43, 35, 25 and 43 genes were specifically upregulated in root cap under C, 1B, 2B, 3B and 5B, respectively (Figure 3C; Supplemental Dataset S11). Additionally, in this cluster, 46 and 11 genes were commonly upregulated between 3B and 5B, and between 1B and 2B, respectively, and also 11 genes were commonly upregulated in 2B, 3B and 5B (Figure 3C; Supplemental Dataset S11). On the other hand, 14, 3, 63, 6 and 4 genes were specifically downregulated in root cap under C, 1B, 2B, 3B and 5B, respectively (Figure 3D; Supplemental Dataset S11). Moreover, 4 genes were commonly upregulated between 3B and 5B, and 3 genes were commonly upregulated in 1B, 2B and 3B, and 3 genes were commonly upregulated in 1B, 2B, 3B and 5B (Figure 3D; Supplemental Dataset S11). 88, 34, 106, 27 and 72 genes were specifically upregulated in QC under C, 1B, 2B, 3B and 5B, respectively (Figure 3E; Supplemental Dataset S11). Additionally, in this cluster, 40 and 8 genes were commonly upregulated between 2B and 3B, and between 1B and 5B, respectively, and also 5 genes were commonly upregulated in 2B, 3B and 5B (Figure 3E; Supplemental Dataset S11). On the other hand, 28, 32, 6 and 9 genes were specifically downregulated in C, 2B, 3B and 5B, respectively (Figure 3F; Supplemental Dataset S11). Also, 9 genes were commonly downregulated between 2B and 3B (Figure 3F; Supplemental Dataset S11). 64, 13, 48 and 37 genes were specifically upregulated in columella under C, 1B, 2B and 5B, respectively (Figure 3G; Supplemental Dataset S11). Moreover, in this cluster, 27 and 6 genes were commonly upregulated between 1B and 2B, and between 2B and 5B, respectively, and also 7 genes were commonly upregulated in 1B, 2B and 5B (Figure 3G; Supplemental Dataset S11). On the other hand, 9, 1, 29 and 9 genes were specifically downregulated in C, 1B, 2B and 5B, respectively (Figure 3H; Supplemental Dataset S11).

### Gene ontology (GO) and KEGG (Kyoto encyclopedia of genes and genomes pathway) orthology (KO) analyses

To determine whether B toxicity is associated with unique GO terms (biological process, cellular component, and molecular function) at different cell clusters, an enrichment analysis of gene expression subsets based on each cluster of B toxicity treatment groups was performed (Supplemental Dataset S12). Accordingly, under 1B condition in columella, top-ranked biological processes were “response to hypoxia, cellular response to hypoxia, response to oxygen level and cellular response to decreased oxygen level”, and top-ranked cellular components were “cell wall and external encapsulating structure”, and top-ranked molecular functions were “serine type endopeptidase inhibitor activity, peptidase inhibitor activity, endopeptidase inhibitor activity, endopeptidase regulator activity, peptidase regulator activity and flavin adenine dinucleotide binding” for upregulated genes (Supplemental Dataset S12). In this cluster, top-ranked biological processes were “tryptophan catabolic process to kynurenine, kynurenine metabolic process, indolalkylamine catabolic process, cellular biogenic amine catabolic process, amine catabolic process and indole-containing compound catabolic process”, and top-ranked cellular component was “mitochondrion”, and top-ranked molecular function was “RNA binding” for downregulated genes (Supplemental Dataset S12). On the other hand, in endodermis, top-ranked biological process was “ATP metabolic process”, top-ranked cellular components were “inner mitochondrial membrane protein complex, mitochondrial protein-containing complex and mitochondrial inner membrane”, and top-ranked molecular function was “protein transmembrane transporter activity” for upregulated genes (Supplemental Dataset S12). In this cluster, top-ranked biological processes were “organonitrogen compound biosynthetic process, cellular amide metabolic process and amide biosynthetic process”, and top-ranked cellular component was “mitochondrion”, and top-ranked molecular function was “FMN binding” for downregulated genes (Supplemental Dataset S12).

In QC, top-ranked biological processes were “electron transport chain, ATP metabolic process and respiratory electron transport chain”, and top-ranked cellular component was “mitochondrion” and top-ranked molecular function was “oxidoreduction-driven active transmembrane transporter activity” for up regulated genes (Supplemental Dataset S12). In root cap, top-ranked biological process was “response to oxygen containing component”, top-ranked cellular component was “anchored component of membrane” and top-ranked molecular function was “mRNA binding” for up regulated genes (Supplemental Dataset S12). In this cluster, top-ranked biological processes were “peptide metabolic process and cellular amide metabolic process”, and top-ranked cellular component was “mitochondrion”, and top-ranked molecular functions were “protein transmembrane transporter activity, glutathione transferase activity and oxidoreduction-driven active transmembrane transporter activity” for downregulated genes (Supplemental Dataset S12).

Moreover, under 2B condition in columella, top-ranked biological processes were “response to hypoxia, cellular response to hypoxia, response to oxygen level and cellular response to decreased oxygen level”, and top-ranked cellular components were “cell wall and external encapsulating structure”, and top-ranked molecular functions were “serine type endopeptidase inhibitor activity, peptidase inhibitor activity, endopeptidase inhibitor activity, endopeptidase regulator activity and peptidase regulator activity” for upregulated genes (Supplemental Dataset S12), In this cluster, top-ranked biological processes were “response to hypoxia, cellular response to hypoxia, response to oxygen level and cellular response to decreased oxygen level”,and top-ranked cellular components were “cell-cell junction, plasmodesma, anchoring junction, symplast and cell junction”, and top-ranked molecular function was “copper ion binding” for downregulated genes (Supplemental Dataset S12). On the other hand, in endodermis, top-ranked biological processes were “response to metal ion, response to inorganic substance and response to cadmium ion”, and top-ranked cellular components were “cell-cell junction, plasmodesma, anchoring junction, symplast and cell junction”, and top-ranked molecular function was “copper ion binding” for upregulated genes (Supplemental Dataset S12). In this cluster, top-ranked biological processes were “response to hypoxia, cellular response to hypoxia, response to oxygen level and cellular response to decreased oxygen level”, and top-ranked cellular component was “vacuole”, and top-ranked molecular function was “modified amino acid binding” for downregulated genes (Supplemental Dataset S12). In QC, top-ranked biological processes were “response to hypoxia, cellular response to hypoxia, response to oxygen level and cellular response to decreased oxygen level”, and top-ranked cellular components were “cell-cell junction, plasmodesma, anchoring junction, symplast and cell junction”, and top-ranked molecular functions were “glutathione transferase activity and oxidoreductase activity” for upregulated genes (Supplemental Dataset S12). In this cluster, top-ranked biological processes were “response to inorganic substance, response to metal ion and response to cadmium ion”, and top-ranked cellular components were “vacuole” and mitochondrion”, and top-ranked molecular functions were “protein tag and ubiquitin protein ligase binding” for downregulated genes (Supplemental Dataset S12). In root cap, top-ranked biological processes were “response to hypoxia, cellular response to hypoxia, response to oxygen level and cellular response to decreased oxygen level”, and top-ranked cellular components were “protein-transporting ATP synthase complex and protein-transporting two sector ATPase complex”, and top-ranked molecular function was “ligase activity” for upregulated genes (Supplemental Dataset S12). In this cluster, top-ranked biological processes were “response to hypoxia, cellular response to hypoxia, response to oxygen level and cellular response to decreased oxygen level”, and top-ranked cellular components were “mitochondrion and vacuole”, and top-ranked molecular functions were “glutathione transferase activity and oxidoreductase activity” for downregulated genes (Supplemental Dataset S12).

Furthermore, under 3B condition in endodermis top-ranked biological processes were “response to inorganic substance and response to cadmium ion”, and top-ranked cellular components were “cell-cell junction, plasmodesma, anchoring junction, symplast, cell junction and vacuole”, and top-ranked molecular function was “copper ion binding” for upregulated genes (Supplemental Dataset S12). In this cluster, top-ranked biological processes were “amide biosynthetic process, peptide metabolic process, cellular amide metabolic process”, and top-ranked cellular component was “ribosome”, and top-ranked molecular functions were “RNA binding, mRNA binding and structural constituent of ribosome” in downregulated genes (Supplemental Dataset S12). In QC, top-ranked biological processes were “response to hypoxia, cellular response to hypoxia, response to oxygen level and cellular response to decreased oxygen level”, and top-ranked cellular components were “cell-cell junction, plasmodesma, anchoring junction, symplast, cell junction, vacuole, cell wall and external encapsulating structure”, and top-ranked molecular function was “modified amino acid binding” for upregulated genes (Supplemental Dataset S12). In this cluster, top-ranked biological process was “response to cadmium ion”, and top-ranked cellular components were “cell-cell junction, plasmodesma, anchoring junction, symplast, cell junction, plastid stroma and chloroplast stroma”, and top-ranked molecular functions were “mRNA binding, copper ion binding and RNA binding” for downregulated genes (Supplemental Dataset S12). In root cap, top-ranked biological processes were “cellular amide metabolic process, amide biosynthetic process, peptide biosynthetic process, peptide metabolic process and translation”, and top-ranked cellular components were “ribosome, ribosomal subunit and cytosolic ribosome”, and top-ranked molecular functions were “structural constituent of ribosome and structural molecule activity” for upregulated genes (Supplemental Dataset S12). In this cluster, top-ranked biological processes were “cellular amide metabolic process, amide biosynthetic process, peptide biosynthetic process, peptide metabolic process, organonitrogen compound biosynthetic process and translation”, and top-ranked cellular component was “ribosome” and top-ranked molecular functions were “structural constituent of ribosome, structural molecule activity, mRNA binding and RNA binding” for downregulated genes (Supplemental Dataset S12).

Also, under 5B condition, in columella, top-ranked biological processes were “response to hypoxia, cellular response to hypoxia, response to oxygen level and cellular response to decreased oxygen level”, and top-ranked cellular components were “cell-cell junction, plasmodesma, anchoring junction, symplast and cell junction”, and top-ranked molecular function was “copper ion binding” for upregulated genes (Supplemental Dataset S12). In this cluster, top-ranked biological process was “response to cadmium ion”, and top-ranked cellular components were “cell-cell junction, plasmodesma, anchoring junction, symplast, cell junction and vacuole”, and top-ranked molecular functions were mRNA binding and RNA binding” for downregulated genes (Supplemental Dataset S12). On the other hand, in endodermis, top-ranked biological processes were “peptide biosynthetic process, amide biosynthetic process, peptide metabolic process, cellular amide metabolic process and translation”, and top-ranked cellular components were “ribosome, ribosomal subunit”, and top-ranked molecular functions were “structural constituent of ribosome and structural molecule activity” for upregulated genes (Supplemental Dataset S12). In this cluster, top-ranked biological processes were “peptide biosynthetic process, amide biosynthetic process, peptide metabolic process, cellular amide metabolic process and translation”, and top-ranked cellular components were “ribosome, ribosomal subunit and cytosolic ribosome”, and top-ranked molecular functions were “mRNA binding, structural molecule activity and structural constituent of ribosome” for downregulated genes (Supplemental Dataset S12). In QC, top-ranked biological process was “ATP metabolic process”, top-ranked cellular components were “inner mitochondrial membrane protein complex and mitochondrial envelope”, and top-ranked molecular functions were “protein transmembrane transporter activity and copper ion binding” for upregulated genes (Supplemental Dataset S12). In this cluster, top-ranked biological processes were “peptide metabolic process and cellular amide metabolic process”, and top-ranked cellular components were “cell-cell junction, plasmodesma, anchoring junction, symplast and cell junction”, and top-ranked molecular function was “protease binding” for downregulated genes (Supplemental Dataset S12). In root cap, top-ranked biological processes were “cellular amide metabolic process, amide biosynthetic process, peptide biosynthetic process, peptide metabolic process and translation”, and top-ranked cellular components were “ribosome, ribosomal subunit and cytosolic ribosome”, and top-ranked molecular functions were “structural constituent of ribosome and structural molecule activity” for upregulated genes (Supplemental Dataset S12). In this cluster, top-ranked biological process was “catabolic process”, and top-ranked cellular components were “cell-cell junction, plasmodesma, anchoring junction, symplast and cell junction”, and top-ranked molecular function was “copper ion binding” for downregulated genes (Supplemental Dataset S12).

To profile specific pathways at cell clusters in response to B toxicity, enrichment analyses of biological pathways defined by KEGG were conducted. KEGG analyses showed that under 1B, in columella, the upregulated DEGs were highly associated with “glutathione metabolism”, “autopathy” and “sulfur metabolism”. In endodermis, the upregulated DEGs were highly associated with pathways including “carbon fixation in photosynthetic organisms”, “glutathione metabolism”, “oxidative phosphorylazation”, “phenylpropanoid biosynthesis”, “glycolysis/gluconeogenesis”, “carbon metabolism” and “biosynthesis of amino acids”, and the downregulated DEGs were highly associated with pathways including “ubiquitin mediated proteolysis”. In root cap, the upregulated DEGs were highly associated with pathways such as “carbon fixation in photosynthetic organisms”, “glycolysis/gluconeogenesis”, “biosynthesis of amino acids”, “cysteine and methionine metabolism” and “carbon metabolism”, and the downregulated DEGs were highly associated with “glutathione metabolism” (Supplemental Dataset S12). Following 2B treatment, in columella, the upregulated DEGs were highly associated with “glutathione metabolism”, “sulfur metabolism” and “alanine, aspartate and glutamate metabolism”. In endodermis, the upregulated DEGs were highly associated “stilbenoid, diaryheptanoid and gingerol biosynthesis”, and the downregulated DEGs were highly associated with pathways including “glutathione metabolism”. In QC, the upregulated DEGs were highly associated with pathways such as “glutathione metabolism”, “phenylpropanoid biosynthesis”, “plant-pathogen interaction” and “MAPK signaling pathway-plant”, and the downregulated DEGs were highly associated with pathways such as “arginine and proline metabolism”, “glutathione metabolism” and “cysteine and methionine metabolism”. In root cap, the upregulated DEGs were highly associated with pathways including “ribosome”, “carbon fixation in photosynthetic organisms”, “glycolysis/gluconeogenesis”, “biosynthesis of amino acids”, “cysteine and methionine metabolism”, “carbon metabolism”, “MAPK signaling pathway-plant” and plant-pathogen interaction”, and the downregulated DEGs were highly associated with pathways including “glutathione metabolism” and “arginine and proline metabolism” (Supplemental Dataset S12). On the other hand, under 3B condition, in endodermis, the upregulated DEGs were highly associated with “glutathione metabolism”, “carbon fixation in photosynthetic organisms”, “glycolysis/gluconeogenesis”, “biosynthesis of amino acids” and “carbon metabolism” and “protein processing in endoplasmic reticulum”, and the downregulated DEGs were highly associated with “oxidative phosphorylation”, “ribosome” and “endocytosis”. In QC, the upregulated DEGs were highly associated with pathways such as “plant-pathogen interaction”, and the downregulated DEGs were highly associated with pathways such as “selenocompound metabolism”, “ribosome”, “biosynthesis of amino acids” and “cysteine and methionine”. In root cap, the upregulated DEGs were highly associated with pathways “ribosome”, “carbon fixation in photosynthetic organisms”, “glycolysis/gluconeogenesis”, “cyanoamino acid metabolism”, “glutathione metabolism”, “biosynthesis of amino acids” and “carbon metabolism”, and the downregulated DEGs were highly associated with “ribosome” (Supplemental Dataset S12). Moreover, under 5B condition, in columella, the upregulated DEGs were highly associated with pathways such as “fructose and mannose metabolism”, carbon fixation in photosynthetic organisms”, “β alanine metabolism”, “alanine, aspartate and glutamate metabolism”, “glutathione metabolism”, “arginine and proline metabolism”, “glycolysis/gluconeogenesis”, “cysteine and methionine metabolism”, “carbon metabolism”, “citrate cycle (TCA cycle)”, “pyruvate metabolism”, “starch and sucrose metabolism” and “biosynthesis of amino acids”, and the downregulated DEGs were highly associated with pathways such as “selenocompound metabolism”, “cysteine and methionine metabolism”, “endocytosis” and “biosynthesis of amino acids”. In endodermis, the upregulated DEGs were highly associated with “ribosome”, “RNA polymerase” and “oxidative phosphorylation”, and the downregulated DEGs were highly associated with pathways such as “selenocompound metabolism”, “ribosome”, “carbon fixation in photosynthetic organisms”, “glutathione metabolism”, “glycolysis/gluconeogenesis”, “endocytosis” and “biosynthesis of amino acid”. In QC, the upregulated DEGs were highly associated with pathways such as “oxidative phosphorylation”, “carbon fixation in photosynthetic organisms”, “flavonoid biosynthesis”, “glycolysis/gluconeogenesis”, “cysteine and methionine metabolism”, “biosynthesis of amino acids”, “alanine, aspartate and glutamate metabolism”, “phagosome”, “glutathione metabolism”, “carbon metabolism” and “phenylpropanoid biosynthesis”, and the downregulated DEGs were highly associated with “glutathione metabolism” and “endocytosis”. In root cap, the upregulated DEGs were highly associated with “ribosome”, and the downregulated DEGs were highly associated with pathways such as “carbon fixation in photosynthetic organisms”, “glycolysis/gluconeogenesis”, “glutathione metabolism”, “biosynthesis of amino acids” and “carbon metabolism” (Supplemental Dataset S12).

As indicated above, glutathonine metabolism was activated caused by B toxicity at different cell clusters in Arabidopsis root. Therefore, we carefully determined the genes related the glutathonine metabolism and they were shown in Table 1. Accordingly, under 1B condition, most significantly upregulated genes were *CICDH* and *GSTF10* in root cap, *GSTU25*, *GSTU24* and *GSTU7* in columella, and *GSTU26*, *GSTF10* and *GPX2* in endodermis. On the other hand, most significantly downregulated genes were *GSTU25*, *GSTU19* and *GSTF8* in root cap (Table 1). Under 2B condition, most significantly upregulated genes were *GSTU25*, *GPX2* and *GSTU5* in columella, *APX1*, *CICDH* and *GSTF10* in endodermis, and *GSTF2*, *GSTU17* and *GSTU11* in QC. On the other hand, most significantly downregulated genes were *GSTU25*, *GSTF8* and *GPX6* in root cap, *GSTF8*, *GSTU24* and *GSTU25* in endodermis, *GSTU24*, *GSTU25* and *GSTU19* in QC (Table 1). Under 3B condition, most significantly upregulated genes were GSTU25, GSTU19 and GSTF8 in root cap, *GSTF4*, *GSTF9* and *GSTF10* in endodermis and *GSTU19*, *GSTU5* and *GSTF8* in QC. On the other hand, *APX1* was most significantly downregulated genes in QC (Table 1). Under 5B condition, most significantly upregulated genes were *GSTU25*, *GSTU22* and *GSTU19* in columella, and *APX1*, *CICDH* and *GSTF10* in QC. On the other hand, most significantly downregulated genes were *APX1*, *GSTU25* and *GSTF8* in root cap, *APX1* in columella, *APX1* and *GSTF8* in endodermis, and *GSTU25* and *GSTU19* in QC (Table 1).

**Table 1.**
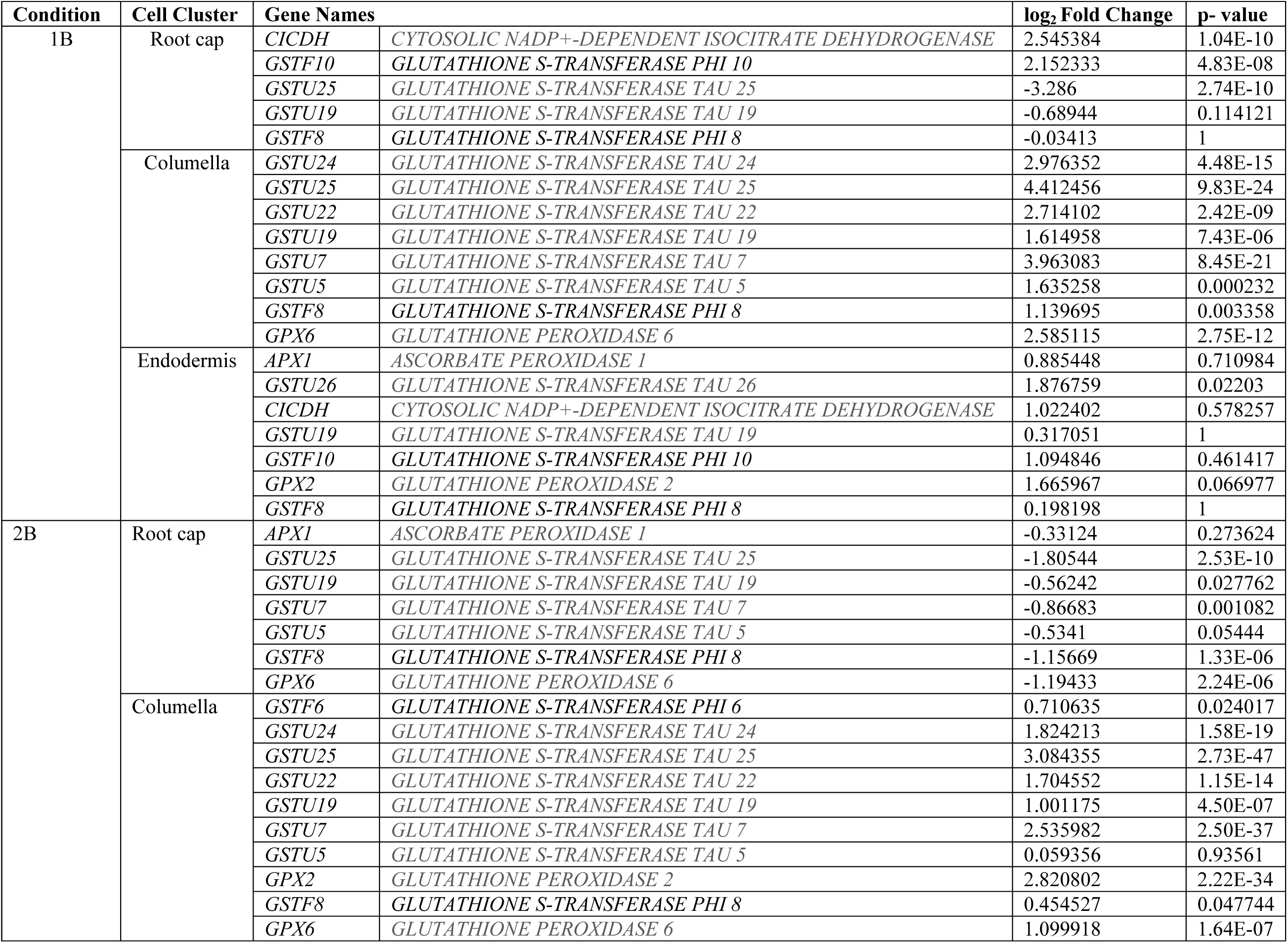
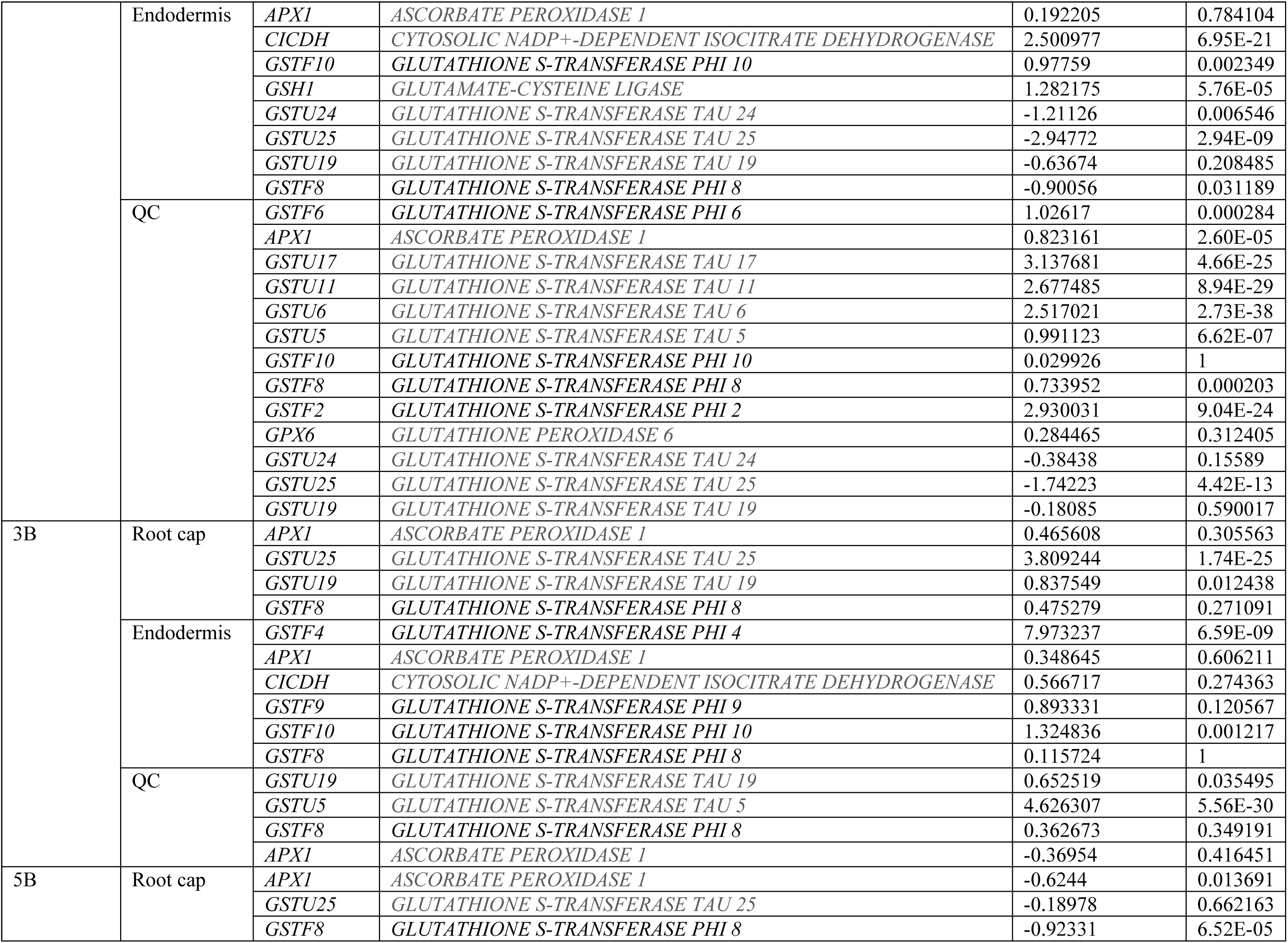
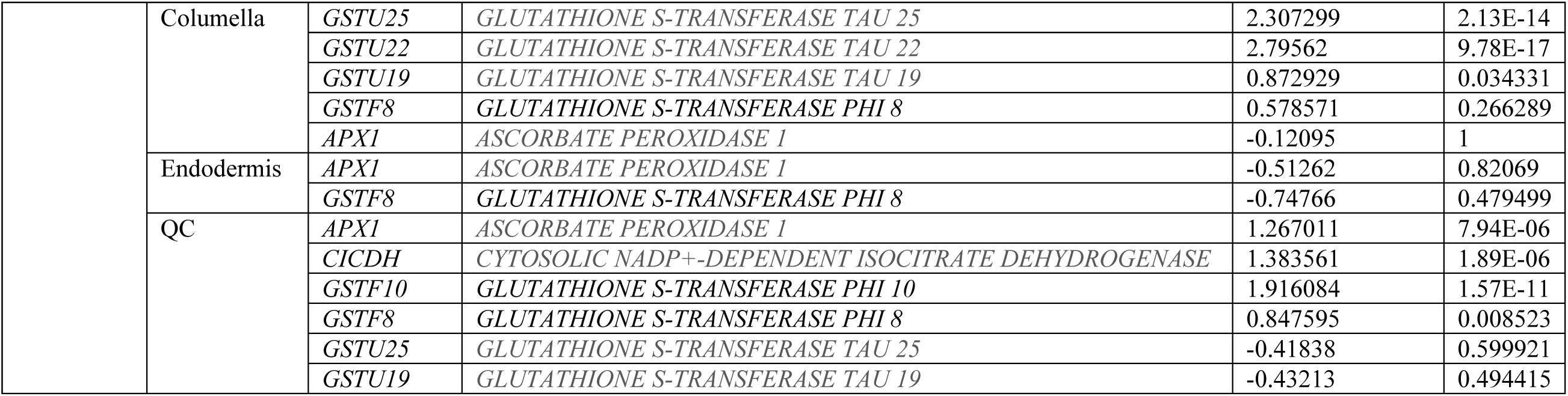
The genes related to glutathione metabolism at different cell clusters under B toxicity in roots of Arabidopsis thaliana

Finally, we identified the transcription factors (TFs) at cell clusters in Arabidopsis root exposed to B toxicity (Supplemental Dataset S13). Accordingly, 13 different TF families including ERF, NAC, C2H2, WRKY, NF-X1, Trihelix, bZIP, bHLH, MYB, C3H, HD-ZIP, LBD and HSF were determined at different cell clusters under B toxicity conditions (Table 2). Under 1B condition, in columella total 14 TFs belonging to 6 TF families such as ERF, NAC, C2H2, WRKY, NF-X1, and Trihelix families were upregulated. In this cluster, the most significantly upregulated TFs were *ANAC087* and *NFXL1* (Table 2). Under 2B condition, in columella, total 14 TFs belonging to 7 TF families including NAC, ERF, LBD, WRKY, bZIP, NF-X1 and Trihelix were upregulated. The most significantly upregulated TFs were *NAC15* and *NAC083* (Table 2). On the other hand, under this condition, in QC, total 33 TFs belonging to 11 TF families such as C2H2, ERF, bHLH, NAC, WRKY, HSF, MYB, C3H, WRKY, bZIP and HD-ZIP were upregulated. The most significantly upregulated TFs were *MYB15* and *MYB108* in QC (Table 2). In root cap, total 12 TFs belonging to 6 TF families including ERF, bHLH, C3H, NAC, WRKY and bZIP were upregulated. The most significantly upregulated TFs were *ERF59* and *ERF109* TFs in root cap (Table 2). Under 3B condition, in QC, total 19 TFs belonging to 9 TF families such as C2H2, ERF, bHLH, NAC, WRKY, HSF, MYB, C3H and bZIP were upregulated. The most significantly upregulated TFs were *MYB15*, *WRKY18* and *WRKY40* in QC (Table 2). Under 5B condition, in columella, 7 TFs belonging to 4 TF families including NAC, C2H2, ERF and bZIP. The most significantly upregulated TFs were *ERF55*, *ANAC081* and *ERF6* in columella (Table 2; Supplemental Dataset S13).

**Table 2.**
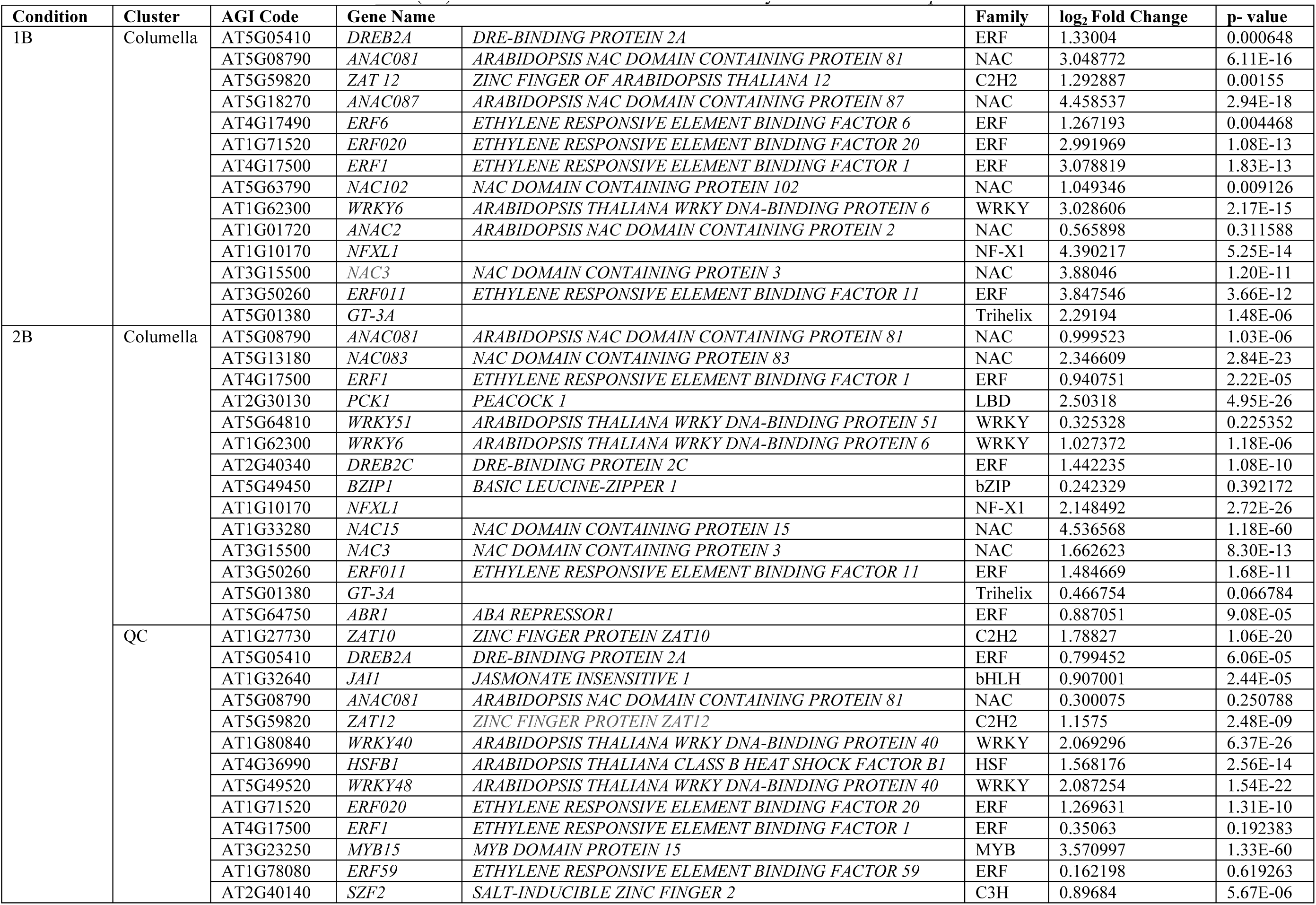
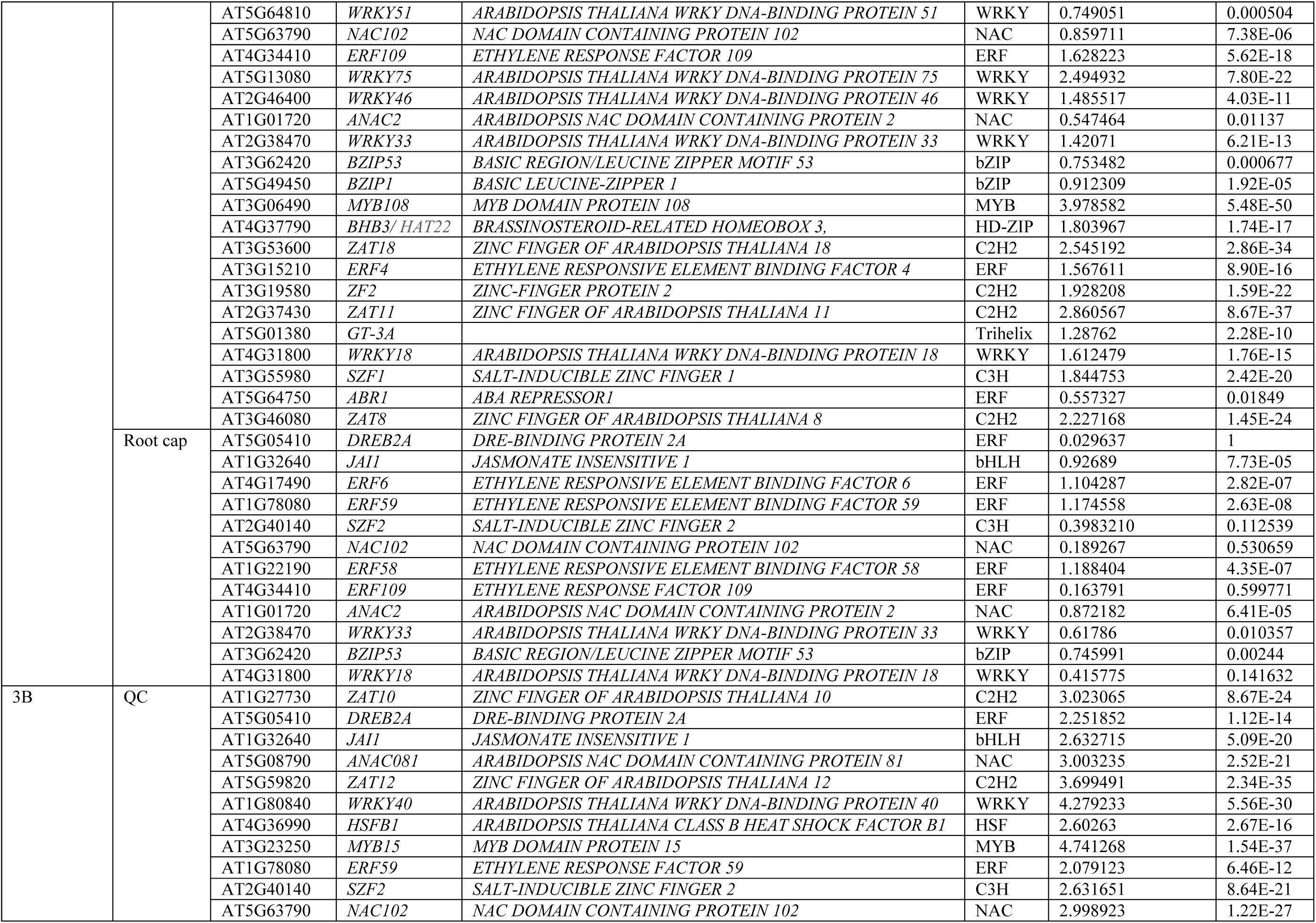
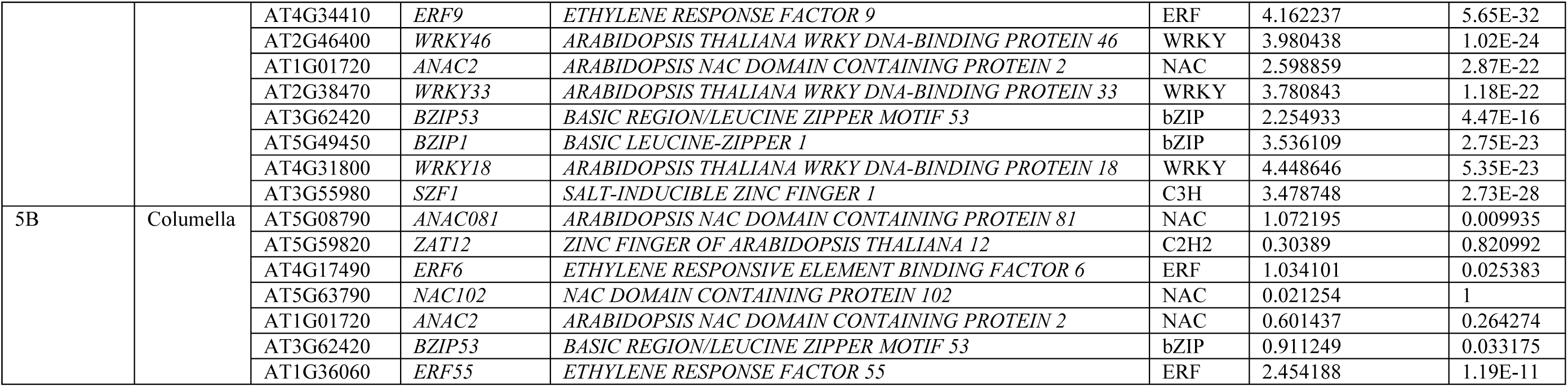
The genes related to Transcription Factor (TF) at different cell clusters under B toxicity in roots of Arabidopsis thaliana

### Heatmap analysis of gene expression

We analyzed the DEGs via heatmap to visualize and interpret the gene expression data at different cell clusters in root tissues of *Arabidopsis thaliana* exposed to B toxicity (Figure 4, A-E). Accordingly, under 1B condition, most significantly upregulated genes were *AT1G12080*, *AT4G22212* and *PDF2.3* in root cap, *AMC9* and *CEL3* in columella, *PME2*, *AIR1B* and *PER57* in endodermis, and *RPS7, RRN26* and *NAD2B* (Figure 4B; Table 3). Under 2B condition, most significantly upregulated genes were *AGP31*, *DFC* and *RBG7* in root cap, *AT3G61930*, *PLP2* and *GLP9* in columella, *PER64*, *DIR9* and *AT1G71740* in endodermis, and *SCREW2*, *VBF* and *PP2B13* in QC (Figure 4C; Table 3). Under 3B condition, most significantly upregulated genes were *GLL23*, *AT2G43610* and *LTP5* in root cap, *AT1G69325*, *AT2G27140* and *RTNLB14* in endodermis, and *PSK2*, *CYP706A2* and *ATBBE8* (Figure 4D; Table 3). Under 5B condition, most significantly upregulated genes were *ORF110B*, *DI21* and *GRP5* in root cap, *JAL9*, *PAC2* and *AT5G60530* in columella, *AT2G07773*, *ATMG00090* and *RPL2* in endodermis, and *EXT17*, *AGP24* and *PER45* (Figure 4E and Table 3). Moreover, *RCPG* was significantly upregulated in columella under 1B and 2B conditions (Figure 4, B-C).

**Figure 4.**
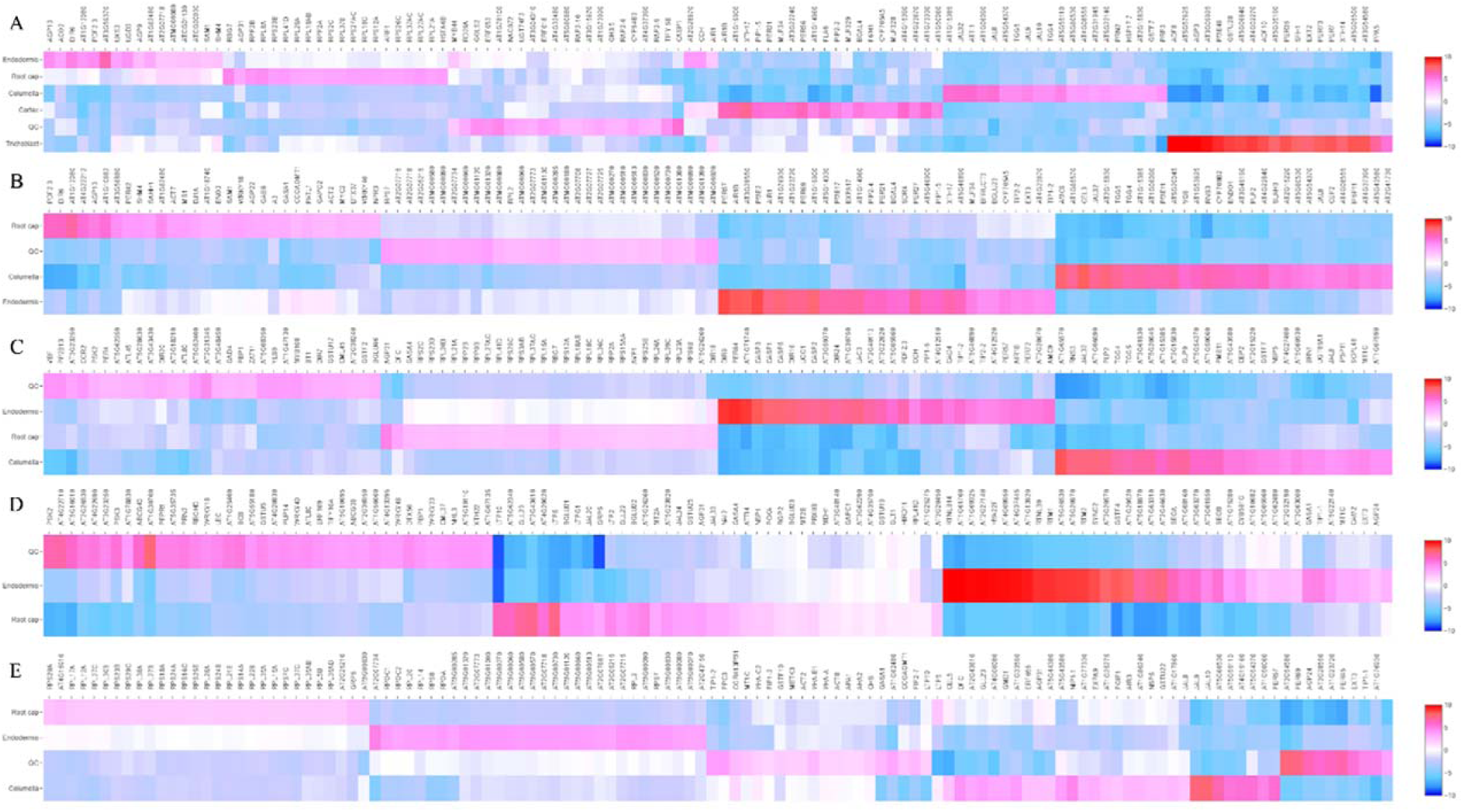
Heatmap visualization of the 50 most differentially expressed genes for each group A) Control group B) 1 mM B treatment group C) 2 mM B group D) 3 mM B group E) 5 mM B group. Red represents up-regulated genes, blue represents down-regulated genes and grey represents unchanged expression.

**Table 3.**
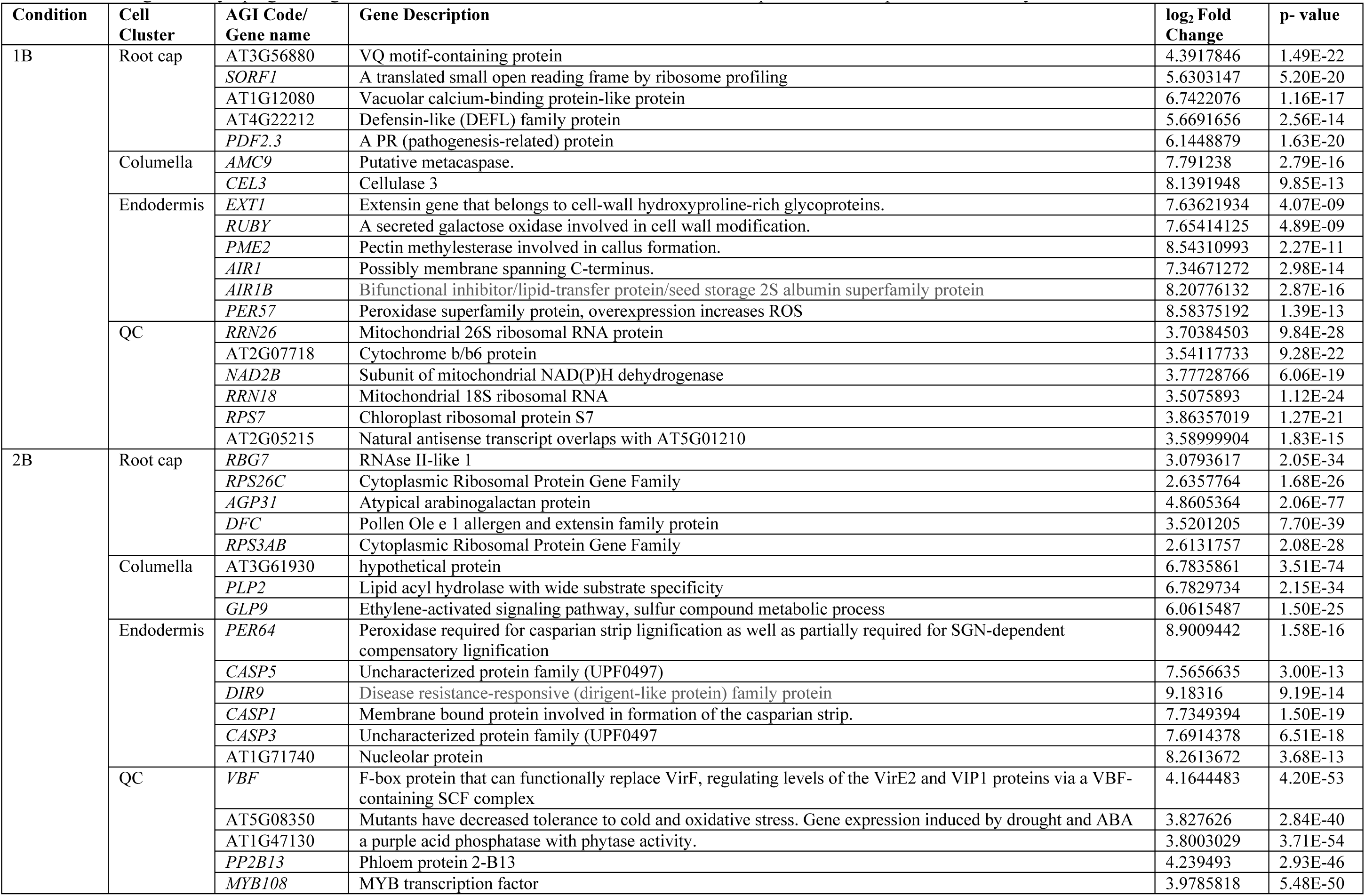
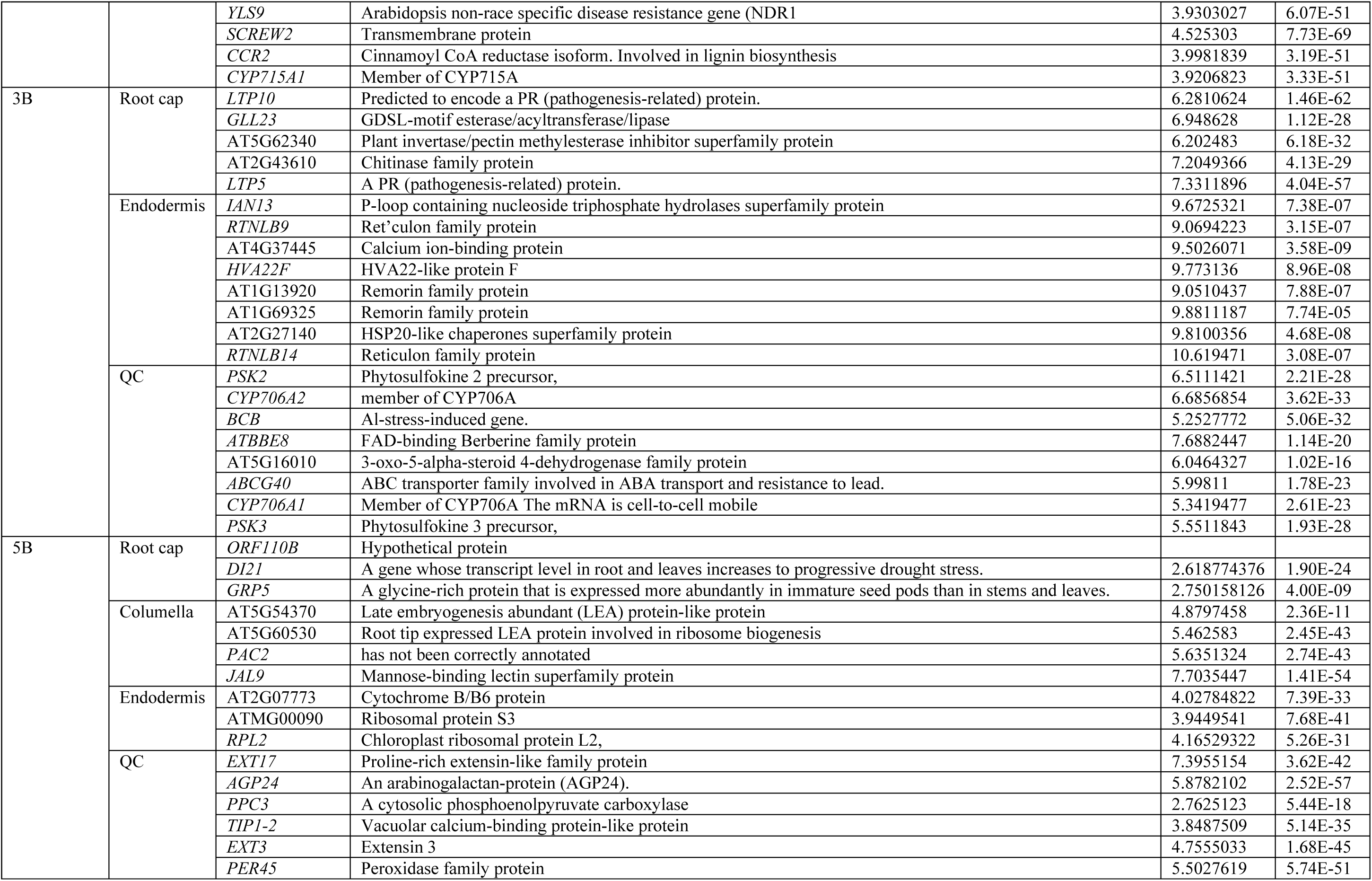
Most significantly upregulated genes at different cell cluster in root tissues of *Arabidopsis thaliana* exposed to B toxicity

## Discussion

B toxicity causes deterioration in developmental and metabolic activities of plants, and thereby yield and serious economic losses (Mengel & Kirkby, 2012). Several transcriptomic studies have been conducted in plants by using microarray and RNA sequencing, which have aimed to find the toxic B responsive regulations at molecular levels (Feng, et al., 2021; Song, et al., 2021; Kayıhan C., 2021; Chen, et al., 2022; Pandey, et al., 2022; Wu, et al., 2022). However, bulk methods are limited in detecting differentially expressed genes at different cell types (Bawa et al., 2022). In addition, it is not possible to detect rare cell types with these methods. Fortunately, scRNA-seq make it possible to profile gene expressions on a cell basis, solve the problem of heterogeneity, and enable analysis of cell types and cell responses to environmental conditions with high resolution and output. Therefore, in this study, a scRNA-seq analysis was performed for Arabidopsis roots exposed to B toxicity at seedling stages to elucidate the molecular basis of the B tolerance mechanism at single cell resolution.

To implement 10X Genomics as a reliable scRNA-seq method, we adapted the established Arabidopsis root protoplasting protocols to produce debris-free single-cell suspensions. These modifications enabled us to successfully generate high-resolution and highly reproducible single-cell transcriptomic maps of 2755 Arabidopsis root cells exposed to B toxicity treatments at seedling stages. Until now, in the studies foucing on scRNA-seq in plants, different numbers of clusters ranging from few to high were found depending on the method (10X Chromium or Drop-seq) and the parameters applied especially for the downstream analysis (Zhang, et al., 2019). Our approach has resulted in the identification of six clusters from the primary root representing some major cell types including endodermis, QC, root cap, columella, cortex, and trichoblast. Endodermis, QC, root cap, columella, cortex and trichoblast were identified in the control groups. In B toxicity treatment groups, endodermis, QC, root cap and columella were identified, however, trichoblast cell cluster was not identified in these groups. This result was compatible with the phenotypic view of Arabidopsis root exposed to all B toxicity conditions because toxic B inhibited the formation of root hair in Arabidopsis.

Similar to previously published scRNA-seq datasets (Shaw et al., 2021), in this study, cell clusters with unknown identity were reported in Arabidopsis root. This might be related to incomplete knowledge of the cell type transcriptomic profiles or a product of technical or computational artifacts or both, such as the presence of two different cells in one droplet barcoded by one bead or RNA leakage during droplet formation (Shaw et al., 2021). Recently, Shaw, et al., (2021) emphasized the issue that existing scRNA-seq bioinformatic pipelines lack functions to address the large cell-size variability and whether this can cause bias in the quantification of transcriptional activity. Despite this situation, unknown clusters, as well as clusters with interesting transcriptomic profiles, can lead to the identification of new cell types and facilitate the in-depth characterization of neglected cell types, respectively (Shaw et al., 2021). One such particular examples were found with cortex cluster of the pooled only in the control group and the absence of columella cluster in 3B. The presence of specific cluster characterized by specific transcripts encoding proteins involved in defense mechanism of plants might present an opportunity to further study the identity and function of this predicted cell type We identified the changes in gene expression profiles in Arabidopsis roots under B toxicity for each cell cluster (Figure 2-3). Accordingly, the number of most significantly upregulated genes under 1B condition was determined in columella (Figure 2). However, they were seen in endodermis under 2B condition (Figure 2G), In addition, the number of most significantly upregulated genes under 3B and 5B conditions were determined in root cap (Figure 2, I-K), This results showed that root cap might have a critical role against severe B toxicity conditions.

To determine and classify functions of DEGs, we performed GO enrichment on the complete set of DEGs using ShinyGO (v.0.76.3) web tool. Analyzing the enrichment of functional categories within the clusters enabled us to perform deeper functional discoveries (Supplemental Dataset S12). Accordingly, in columella, in the category of molecular functions, “serine type endopeptidase inhibitor activity”, “peptidase inhibitor activity”, “endopeptidase inhibitor activity” and “peptidase regulator activity” were the top enriched GO terms among upregulated genes of B treatment conditions. Likewise, it was found that the well-represented molecular functions were peptidase and endopeptidase inhibitor activity for upregulated genes in roots of two contrasting wheat cultivars (Kayıhan et al., 2017). On the other hand, in root cap, in the category of cellular component, “ribosome”, “ribosomal subunit” and “cytosolic ribosome”, and in the category of molecular functions, “structural molecule activity” and “structural constituent of ribosome” were the top enriched GO terms among upregulated genes of B treatment conditions. Similarly, remarkable number of genes related to ribosomal proteins was significantly regulated in wheat cultivars following B-toxicity (Kayıhan et al., 2017).

In order to know which metabolic pathways were affected under B toxicity, all DEGs were analyzed with ShinyGO (v.0.76.3) for KEGG analysis (Supplemental Dataset S12). Accordingly, for all cell clusters, under B toxicity conditions, the DEGs were significantly enriched in 33 KEGG pathways, mainly such as “glutathione metabolism”, “autopathy”, “sulfur metabolism”, “carbon fixation in photosynthetic organisms”, “cysteine and methionine metabolism”, “carbon metabolism”, “citrate cycle (TCA cycle)”, “pyruvate metabolism”, “starch and sucrose metabolism”, “biosynthesis of amino acids”, “phenylpropanoid biosynthesis”, “plant-pathogen interaction”, “MAPK signaling pathway-plant”, “flavonoid biosynthesis, “ribosome”, “ubiquitin mediated proteolysis”, “arginine and proline metabolism”, “endocytosis”, (Supplemental Dataset S12). Likewise, it has been recently shown that biosynthesis of secondary metabolites, plant– pathogen interaction, metabolic pathways, phenylpropanoid biosynthesis, RNA transport, and MAPK signaling pathway (plant) are the pathways with the highest number of DEGs in *Triticum zhukovskyi* under B toxicity (Pandey, et al., 2022). Furthermore, the importance of phenylpropanoid pathways has been shown to play a role in the compartmentalization of B in vacuoles in *Arabidopsis thaliana* (Kayıhan et al., 2021). Also, Kayıhan et al., (2017) found that the majority of differentially expressed genes related to protein metabolism were involved in protein degradation in response to B toxicity in sensitive and tolerant wheat cultivars and in fact, the numbers of these genes were higher in root tissues of sensitive wheat cultivars than tolerant wheat cultivar under B toxicity.

In columella, endodermis and QC under B toxicity conditions, upregulated genes were highly associated with “glutathione and “sulfur metabolism” (Supplemental Dataset S12). It has been suggested that B-anthocyanin complexes in vacuoles are an internal mechanism of tolerance to B toxicity (Landi, et al., 2015). Anthocyanin–glutathione or – glutathione –S transferase (GST) complexes can temporarily bind to metal or metalloid ions. In this way, GST-anthocyanin-metal complexes are formed and/or glutathionylanthocyanin metal complexes are vacuolated sequestered (Landi, et al., 2015). Furthermore, GST-anthocyanin-metal complexes may be exported by ABC transporters. Particularly, in columella, *GSTU24, GSTU25*, *GSTU22*, *GSTU19*, *GSTU7*, *GSTU5*, *GSTF8* and *GPX6* under 1B condition, *GSTF6, GSTU24, GSTU25, GSTU22, GSTU19, GSTU7, GSTU5, GPX2, GSTF8 and GPX6,* under 2B condition, and *GSTU25*, *GSTU22*, *GSTU19*, *GSTF8* and *APX1* under 5B condition were revealed to be enriched among upregulated genes in functions associated with glutathione metabolism. Additionally, in Columella, AT1G55920, AT4G04610, AT4G21990 under 1B condition, and AT1G55920 AT1G62180, AT4G04610 and AT4G21990 under 2B condition were revealed to be enriched among upregulated genes in functions associated with “sulfur metabolism”. These results might indicate the importance of GST related to an internal B tolerance mechanism in columella cell cluster in Arabidopsis root. In another study, Kayıhan et al., 2021 examined toxic B-treated *Arabidopsis thaliana* to determine changes in the expression levels of genes involved in anthocyanin biosynthesis and transport, and related transcription factors (TFs). Accordingly, 3 mM boric acid treatment induced expression levels of anthocyanin biosynthesis genes (*4CL3*, *C4H*) and TFs (MYB75 and MYB114) and anthocyanin transporter genes (*TT13* and *TT19*). In this study, *C4H* (AT2G30490) was commonly upregulated between endodermis and QC under 2B condition and significantly upregulated in QC under 3B condition. Also, we found that in QC AT1G14540 AT1G61820 AT1G80820 AT2G30490 AT2G37040 AT4G34230 AT5G39580 under 2B condition, and *PER45*, *CCoAOMT1* and *PER69* under 5B condition were revealed to be enriched among upregulated genes in functions associated with “phenylpropanoid biosynthesis”. Moreover, in plants, cysteine biosynthesis plays a central role in fixing inorganic sulfur from the environment and provides the only metabolic sulfide donor for the generation of methionine, glutathione, iron-sulfur clusters, and multiple secondary metabolites (Bonner, et al., 2005). KEGG analysis showed that in root cap, *SAM1*, *SAMDC1*, *SAHH1* and *MS1* under 1B condition, and *SAT1*, *SAM2*, *SAHH1* and *MS1* under 2B condition were revealed to be enriched among upregulated genes in functions associated with “cysteine and methionine metabolism”. On the other hand, in QC, *TAT3*, *SAMDC1* and *MS1* under 2B condition, and *SAHH1* and *MS1* under 3B condition were revealed to be enriched among downregulated genes in functions associated with “cysteine and methionine metabolism”. These results may indicate that cysteine and methionine metabolism play a key role in the formation of GST-anthocyanin-metal complexes related to the B tolerance mechanism by contributing to sulfur uptake in the root cap and QC.

Photosynthesis is one of the main metabolic processes impaired by B excess, mainly by reducing photosynthetic rate (Pn), chlorophyll content, electron transport rate, CO_2_ assimilation, by impairing photosystem II (PSII) efficiency and by triggering the production of reactive oxygen species in the chloroplast. An approach to alleviating B toxicity in plants is to increase their physiological tolerance, including the enhancement of photosynthesis and the prevention of oxidative stress, by applying nutrient elements, plant growth regulators, vitamins and biostimulators (Antonopoulou & Christos Chatzissavvidis, 2022). KEGG analysis showed that in QC, ATCG00020, ATCG00130, ATCG00340, ATCG00470 and ATCG00720 under 2B condition, and *PSBA*, *PSAB*, *ATPE*, *PETB* under 3B condition were revealed to be enriched among upregulated genes in functions associated with “Photosynthesis” (Supplemental Dataset S12). Moreover, in root cap, *GAPC2*, *GAPC1* and *FBA8* under 1B condition, *GAPC2*, *GAPC1*, *FBA8*, *CTIMC* and *PCKA* under 2B condition, and *GAPC2*, *GAPC1*, *FBA8*, *CTIMC* and *PCKA* under 3B condition were revealed to be enriched among upregulated genes in functions associated with “carbon fixation in photosynthetic organism”. Furthermore, AT1G04410, AT1G13440, AT1G65930, AT3G04120, AT3G14940, AT3G52930, AT3G55440 and AT4G14880 under 1B condition and AT1G13440, AT3G04120, AT3G52930 and AT3G55440 3B condition were revealed to be enriched among upregulated genes in functions associated with “carbon fixation in photosynthetic organisms” (Supplemental Dataset S12). In root cap, *GAPC2*, *GAPC1*, *FBA8* and *MS1* under 1B condition, *GAPC2*, *GAPC1*, *FBA8*, *CTIMC* and *PCKA* under 2B and 3B conditions condition were revealed to be enriched among upregulated genes in functions associated with “carbon metabolism”. Moreover, in endodermis, AT1G13440, AT3G04120, AT3G14940, AT3G52930 and AT3G55440 under 1B condition, and AT1G13440, AT3G04120, AT3G55440 and AT3G52930 under 3B condition were revealed to be enriched among upregulated genes in functions associated with “carbon metabolism” (Supplemental Dataset S12). This might be related to the ability of B to form complexes with molecules such as ATP and NADPH (Cervilla, et al., 2009). This interaction limits the availability of free energy required for carbohydrate biosynthesis in the Calvin–Benson cycle. Thus, alterations in sugar content and partitioning (Roessner, et al., 2006; Papadakis, et al., 2018) and carbon skeleton devoted to amino acids can be observed under B toxicity (Guo, et al., 2014; Sang, et al., 2015). In relation to carbohydrate biosynthesis, in root cap, *GAPC2*, *ENO2*, *GAPC1* and *FBA8* under 1B condition, *GAPC2*, *ENO2*, *GAPC1*, *FBA8*, *CTIMC* and *PCKA* under 2B condition and *GAPC2*, *ENO2*, *GAPC1*, *FBA8*, *CTIMC* and *PCKA* under 3B condition were revealed to be enriched among upregulated genes in functions associated with “glycolysis/gluconeogenesis”. Additionally, in endodermis, AT1G13440, AT3G04120, AT3G52930 and AT3G55440 under 1B condition, and AT1G13440, AT2G36530, AT3G04120, AT3G52930 and AT3G55440 under 3B condition were revealed to be enriched among upregulated genes in functions associated with “glycolysis/gluconeogenesis”.

Induction of jasmonic acid (JA) related genes was found to be an important late response to B toxicity (Öz, et al., 2009). Our study showed that several JA related genes were upregulated under B toxicity. Accordingly, AT3G56880 was specifically upregulated in root cap under 1B and 2B conditions and QC under 3B condition (Figure 2 and 4; Supplemental Dataset S10-11). Furthermore, *AGP31* was specifically upregulated in root cap under 1B, 2B and 3B conditions and commonly upregulated between columella and root cap under 5B condition (Figure 2 and 4; Supplemental Dataset S10-11). *NOI5* (AT3G48450) was specifically upregulated in QC under 2B condition. PSK2 (AT2G22860) was specifically upregulated in QC under 2B and 3B conditions (Figure 2 and 4; Supplemental Dataset S10-11). *ABCG40* (AT1G15520) was specifically upregulated in columella under 1B and 2B conditions and in QC under 5B condition (Figure 2 and 4; Supplemental Dataset S10-11).

TFs are multi-functional proteins that may simultaneously control numerous pathways during stresses in plants. In recent years, numerous plant TFs involved in B toxicity have been identified including WRKY, ERF, NAC, MYB (Aquea, et al., 2012; Tombuloglu, et al., 2013; Kayıhan, et al., 2017; Hua, et al., 2017; Chen, et al., 2022). In our study, we identified the 13 different TF families including ERF, NAC, C2H2, WRKY, NF-X1, Trihelix, bZIP, bHLH, MYB, C3H, HD-ZIP, LBD and HSF at different cell clusters under B toxicity conditions (Table 2). Particularly, *ANAC081*, *ANAC2* and *NAC102* were highly upregulated under B toxicity conditions. Interestingly, ANAC081 was commonly upregulated in columella under all B toxicity conditions (Supplemental Dataset S13). On the other hand, the greatest number of TFs expression was seen in QC. Accordingly, C2H2, ERF, bHLH, NAC, WRKY, HSF, MYB, C3H, bZIP, HD-ZIP and Trihelix TF families were upregulated in QC (Supplemental Dataset S13). *SZF2*, *ZAT12*, *ANAC2*, *NAC102*, *WRKY46*, *JAI1*, *HSFB1*, *SZF1*, *WRKY40*, *WRKY33*, *BZIP1*, *DREB2A*, *MYB15*, *WRKY18*, *ANAC081*, *BZIP53*, *ZAT10* and *ERF59* TFs were commonly upregulated between 2B and 3B (Supplemental Dataset S13). From these results, it can be concluded that QC and columella are highly involved in TF regulation under B toxicity.

In conclusion, we have created a high-resolution single-cell transcriptomic map and identified cell specific transcriptome patterns Arabidopsis root under B toxicity conditions. The pathways already presented in the literature related to B toxicity were found and many new genes specific to cell type were identified. Further analyses of these genes and pathways at cell cluster may be crucial to clarify toxic B responsive regulation and B tolerance mechanism more accurately in plants.

## Materials and methods

### Plant growth and stress treatment

Wild type (WT) *Arabidopsis thaliana* cv. Columbia seeds were provided by Assistant Prof. Dr. Emre Aksoy at Middle East Technical University in Turkey. Before sowing, Arabidopsis seeds were surface sterilized using 70% (v/v) EtOH for 2 minutes and 2.5% (v/v) NaOCl for 10 minutes. After washing three times with distilled water for approximately 10 seconds each, surface sterilized seeds were grown on Murashige and Skoog (MS) growing medium. Briefly, control groups were grown in boric acid-free MS media (pH: 5.7) for 14 days. 1- or 2-mM B toxicity treatment groups were grown in MS media containing 1- and 2-mM boric acid, respectively for 14 days. The 3- and 5-mM toxicity treatment groups were grown in MS medium without boric acid for 8 days, followed by another 6 days in MS medium containing 3- and 5- mM boric acid, respectively. Plates were kept at +4 °C for 3 days and then positioned vertically for root development in the growth chamber. All groups were grown in 3 replicates at 22°C ±1°C with 16 hours of light (200 μmol m^−2^ s^−1^) and 8 hours of dark photocycle at 60% relative humidity.

### Protoplasting

At 14 d after germination, approximately 30 primary roots were chopped with a length of 2 cm, above the tips with the help of forceps and placed into a 35-mm-diameter dish containing a 70 µm strainer and 3 ml enzyme solution (1.25% [w/v] Cellulase [“ONOZUKA” R10, Yakult], 0.1% [w/v] Pectolyase [P-3026, Sigma-Aldrich], 0.4 M Mannitol, 20 mM MES [pH 5.7], 20 mM KCl, 10 mM CaCl2, 0.1% [w/v] bovine serum albumin). Petri dishes was shaken in a shaker at 90 rpm for 2 and half hours, gently crushed every half hour. The liquid parts that filtered out of the petri dish were taken into 15 ml falcon and strained through a 40 µm pipette strainer. The filtered solution was centrifuged at 100 *g* for 6 min at 22°C. Supernatants were gently removed without damaging the pellets. The pelleted protoplasts resuspended in 500 µl of 8% mannitol. The protoplast solution was strained through a 40-pipette strainer to achieve the desired cell concentration (10^4^ protoplasts/ml). 0.4% Tyrphan Blue was added to 1.5 ml eppendorfs containing 25 µl of protoplast solutions. After waiting 1-2 minutes, they were loaded onto a Thoma slide and viewed with a light microscope. Live/dead cells in 1 µl of each sample were counted separately and cell viability was calculated.

### Construction and sequencing of the single-cell RNA-seq libraries

The protoplast suspension was loaded into Chromium microfluidic chips with 3’ v2 chemistry and barcoded with a Chromium Controller (10X Genomics). RNA from the barcoded cells was subsequently reverse-transcribed and sequencing libraries constructed with reagents from a Next GEM Single Cell 3ʹ Reagent Kits v3.1 (10X Genomics) according to the manufacturer”s instructions. Sequencing was performed with Illumina NovaSeq 6000 according to the manufacturer”s instructions (Illumina). The sequencing data are available at the NCBI Gene Expression Omnibus repository under accession number 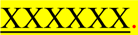.

### Generation and analysis of single-cell transcriptomes

Raw reads generated by Illumina sequencers were demultiplexed into FASTQ files, mapped to the TAIR10 reference genome using STAR software (Spliced Transcripts Alignment to a Reference) and performed filtering, barcode counting and UMI counting by 10X Genomics Cell Ranger pipeline (v3.0.0) using default parameters. The Cell Ranger (v3.0.0) filters and aligns the reads from the file containing the reads for each sample and counts the unique molecular identifiers (UMIs) of the aligned reads to quantify the expression of each gene in each cell. For each gene and each cell barcode (filtered by Cell Ranger (v3.0.0)), unique molecule identifiers were counted to construct digital expression matrices. Finally, datasets were aggregated for use in Loupe Browser (v.6.2.0). aggregation automatically equalizes the average read depth per cell between groups before combining the sample files. This approach made it possible to avoid artifacts that may arise due to differences in sequencing depth.

### Dimensionality reduction, cell clustering and cluster identification

All downstream single-cell analyses were performed using the Loupe Browser (6.2.0) that allows easy visualization and analysis of Chromium single-cell 5′ and 3′ gene expression data, using default parameters unless specifically noted. This software was used to identify significant genes and cell types and differential gene expression. Firstly, interactive filtering and reclustering workflow were used to precisely screen out possible cell multiplets, dead cells, or cells with low diversity and perform Principle Component Analysis (PCA) and t-distributed Stochastic Neighbor Embedding of principal components (t-SNE). In this workflow, filtering was performed using violin plots of UMI counts of the currently selected barcodes, threshold by a distinct number of detected features (number of distinct genes found for each barcode) and the percentage of UMIs per barcode associated with mitochondrial genes. Then, normalization was performed with the library size parameter per cell. PCA (default 20 PCA) was performed via the num_principal_comps command using the Python implementation of the IRLBA algorithm to reduce the size of the dataset, the samples were visualized t-SNE (default 30 t-SNE. After filtering and reclustering workflow, cell clusters were then identified using specific and validated gene markers for each cell type to cluster the cells in a robust manner.

### Differential gene expression of single-cell transcriptome of *Arabidopsis* roots exposed to B toxicity

Differential gene expression between clusters was determined with a negative binomial exact test using sSeq. Clusters were run through the algorithm and compared against all other cells and within clusters to yield genes that were differentially expressed in the cluster relative to the rest of the sample. Cell Ranger (v3.0.0) performed batch effect corrections, which allowed the samples to be run in separate wells and combine the datasets from these runs. The algorithm identified similar cell subpopulations between batches and merged them. The algorithm is based on the “Mutual Nearest neighbor” method (i.e., a pair of cells from 2 different batches that contain each other”s nearest neighbor).

### Gene ontology and KEGG orthology analysis

An online tool, ShinyGO (v.0.76.3) was used for comparing and plotting Gene Ontology (GO) annotation and KEGG annotation results at specific cell clusters (Ge et al., 2020). p-values were calculated according to the hypergeometric distribution of gene numbers. False Discovery Rate (FDR) was calculated according to the nominal p-value obtained from the hypergeometric test. The FDR cut-off value was chosen as 0.05. then the GO terms (biological process, cellular component, and molecular function) and KEGG pathways were ranked according to the FDR Enrichment value and visualized with Dotplot. KEGG pathways were obtained and visualized with KEGG pathway map and Dotplot.

### Heatmap analysis of gene expression

Lists of 50 most significantly differentially expressed genes were used for the heat map analysis. Briefly, gene count was log_2_ normalized and scaled via the Loupe Browser (v.6.2.0). A hierarchical clustering was carried out to cluster both rows and columns using Euclidean distance.

## Author Contributions

C.K. conceived and designed the study and performed the experiments and wrote the manuscript.

H.Y. analyzed the data and wrote the manuscript. H.B.Ü and O.Y. performed the experiments.

E.A. edited the manuscript. All authors read and approved the final manuscript.

## Acknowledgement

We would like to thank Gen Era Diagnostic and Aydan SARAÇ DERDİYOK at 10X Genomics for their contributing efforts to the emergence of this work. No conflict of interest is declared.

